# Vigilance response of a key prey species to anthropogenic and natural threats in Detroit

**DOI:** 10.1101/2020.06.09.142992

**Authors:** Samantha Lima, Siria Gámez, Nathaniel Arringdale, Nyeema C. Harris

## Abstract

Rapid urbanization coupled with increased human activity induces pressures that affect predator-prey relations through a suite of behavioral mechanisms, including alteration of avoidance and coexistence dynamics. Synergisms of natural and anthropogenic threats existing within urban environments exacerbate the necessity for species to differentially modify behavior to each risk. Here, we explore the behavioral response of a key prey species, cottontail rabbits (*Sylvilagus floridanus*), to pressures from humans, domestic dogs, and a natural predator, coyotes (*Canis latrans*) in a human-dominated landscape. We conducted the first camera survey in urban parks throughout Detroit, Michigan in 2017-2020 to assess vigilance response corresponding to a heterogeneous landscape created from variation in the occupancy of threats. We predicted a scaled response where cottontail rabbits would be most vigilant in areas with high coyote activity, moderately vigilant in areas with high domestic dog activity, and the least vigilant in areas of high human activity. From 8,165 independent cottontail rabbit detections in Detroit across 11,616 trap nights, one-third were classified as vigilant. We found vigilance behavior increased with coyote occupancy and in locations with significantly high domestic dog activity, but found no significant impact of human occupancy or their spatial hotspots. We also found little spatial overlap between rabbits and threats, suggesting rabbits invest more in spatial avoidance; thus, less effort is required for vigilance. Our results elucidate strategies of a prey species coping with various risks to advance our understanding of the adaptability of wildlife in urban environments. In order to promote coexistence between people and wildlife in urban greenspaces, we must understand and anticipate the ecological implications of human-induced behavioral modifications.

## Introduction

The 20th and 21st centuries have seen unprecedented population growth and expansion of cities, with 60% of the global population expected to live in urban centers by the year 2030 (United Nations, 2018). Urbanization coupled with other increased anthropogenic pressures has fundamentally changed ecosystems worldwide (Foley et al 2005, Grimm et al 2008, Pickard et al 2017, Chen et al 2020). Cities result in fragmenting natural habitats that restrict gene flow, change species assemblages, and alter the behavior of animals and people alike (Romano 2002, Tigas et al 2002, Crooks et al 2004, Lowry et al 2013, Johnson and Munshi-South 2017). These environmental perturbations have implications for wildlife and a myriad of ecological interactions including predator-prey relationships.

Non-consumptive fear effects induced by humans are pervasive in urban environments and drive behavioral changes in wildlife (Ciuti et al 2012, Gaynor et al 2018). For example, eastern grey squirrels (*Sciurus carolinensis*) in New York City have become sensitive to human movements and show behavioral plasticity in their ability to adjust flight initiation distance based on human activity (Bateman and Fleming 2014). Exposure to human audio cues reduced foraging time and increased the amount of time spent being vigilant in badgers (*Meles meles*) in Great Britain as compared to exposure to non-human predator audio cues (Clinchy et al 2016). Behavioral plasticity in predator and prey species alike directly influence their ability to avoid and coexist with intense human pressures in urban centers (Muhly et al 2011, Lowry et al 2013). While prey modify their behavior to avoid attempted predation, predators modify their behavior to account for prey behavior and to increase the likelihood of success of their predation attempts. Specifically, prey are forced to modify their behavior spatially or temporally to avoid threats from humans as well as associated domestic animals or natural predators (Fenn and Macdonald 1995, Gliwicz et al 2008, Reilly et al 2017). Modification of behavior has therefore become necessary for the survival of both predators and prey in urban environments, as risks govern behavior (Lima 1998). However, despite the recent burgeoning of urban ecology studies, how humans and domestic animals alter mammalian vigilance behavior remains understudied.

Highly adaptable species and those with relatively smaller body sizes are more successful at coexisting with humans in urban areas (Bateman and Flemming 2012). Carnivores, particularly large bodied carnivores, have historically faced intense persecution from humans (Munoz-Fuentes et al. 2010). Large predators depredate livestock and compete with humans for resources including space and prey, often resulting in humans employing lethal interventions (Mech 1995, Witmer and Whittaker 2001, Treves 2003, Muhly and Musiani 2009). However, many mid to small-sized predators are able to thrive in areas of high anthropogenic influence (Wilkinson and Smith 2001, Ikeda et al 2004). In particular, coyotes (*Canis latrans*) have adapted to living with humans in part, by exploiting anthropogenic food subsidies and shifting diurnal movement in response to human disturbance (Kitchen et al 2001, Gese and Bekoff 2004). This, in conjunction with wide extirpations of the grey wolf (*Canis lupus*) has allowed coyotes to expand their range to the entirety of the United States beyond previous restrictions to the central and western portions of the country (Crooks 1999, Hody and Kays 2018). Domestic dogs (*Canis familiaris*) have similarly become abundant within urban areas and thus, can exert top-down pressures as a member of the carnivore community (Ordeñana et al. 2010). These ecological and behavioral changes in carnivores can have cascading effects on their prey species, subsequently altering their behavior.

Concurrent with predators employing strategies for coexistence, their prey must also mitigate risks in human dominated landscapes. Threats for prey species in urban environments are often exacerbated by multiple sources including direct mortality from natural and anthropogenic sources. Prey may employ similar strategies to mitigate risks from humans as they do to mitigate risks from natural predators (Parsons et al. 2016). As such, fear effects in urban environments can result in prey modifying temporal activity or habitat selection to reduce predation risks (Chambers and Dickman 2002, Dowding et al. 2010). Discernment between immediate and distal threats requires delegating time to vigilance in order to assess and respond to risks across the landscape. However, there are tradeoffs because more time spent being vigilant means less time foraging, mating, and performing other behaviors like grooming (Quenette 1990). Environmental conditions including vegetation height, tree cover, and the distribution of water sources can interact to produce varying levels of predation risk and thus influence the amount of time prey spend being vigilant (Scheel 1993, Tchabovsky et al 2001).

Cottontail rabbits (*Sylvilagus floridanus*) are a key prey source for many mammalian carnivores as well as avian predators and occasionally snakes in urban environments throughout the United States (Beasom and Moore 1977, Litvaitis and Shaw 1980, Wittenberg 2012). Because rabbits are an important part of coyotes’ diet, along with small rodents, coyotes exert top-down pressures to control their populations (Poessel, Mock, and Breck 2017). Cottontail rabbits have high reproductive rates that result in rapidly growing populations that interact, directly or indirectly, with humans in gardens, yards, parks and other green spaces throughout city limits (Hunt et al 2014, Baker et al 2015). We conducted a non-invasive camera survey to investigate the vigilance behavior of rabbits in response to anthropogenic and natural threats. Our work occurred throughout Detroit, the largest city in Michigan, located in the Great Lakes region of the USA from 2017-2020. Here, we delineated human, coyote, and domestic dog risk zones to detect differences in cottontail vigilance response and investigated the potential factors influencing vigilance.

Species exploiting urban environments may exhibit higher plasticity to cope and acclimate with anthropogenic threats (Samia *et al.* 2015). The gray squirrel (*Sciurus carolinensis),* another common urban prey species, is less wary of humans in areas more densely populated by humans (Parker and Nilon 2008). This suggests a level of acclimation to human presence, which we reasonably anticipate occurring in cottontail rabbits who are exposed to similar pressures of human activity in an urban environment. Therefore, we expect a similar level of acclimation in cottontail rabbits where they are less vigilant in areas heavily populated by humans. Because of the similarities in body size and behavior between domestic dogs and coyotes, we anticipate rabbits will show more vigilance in areas with high domestic dog presence than areas with high human presence. However, domestic dog populations are generally larger in urban areas because of association with humans. In Detroit, we anticipate some level of acclimation to their presence from cottontail rabbits and therefore, we expect the response to dogs to be less dramatic than the response to coyotes. However, unaccompanied dogs could illicit pronounced fear responses. Overall, we expect a scaled response where rabbits will be least vigilant in areas with high human activity, with vigilance response increasing slightly in the areas with high domestic dog activity, and the most vigilance being displayed in areas of high coyote activity, as coyotes are an actual formidable predator of rabbits (Figure 1). Results will further our understanding of how a key prey species behaves in dynamic urban landscapes, information necessary to foster safe and positive interactions between people and wildlife coexisting in the city.

**Figure 1.**
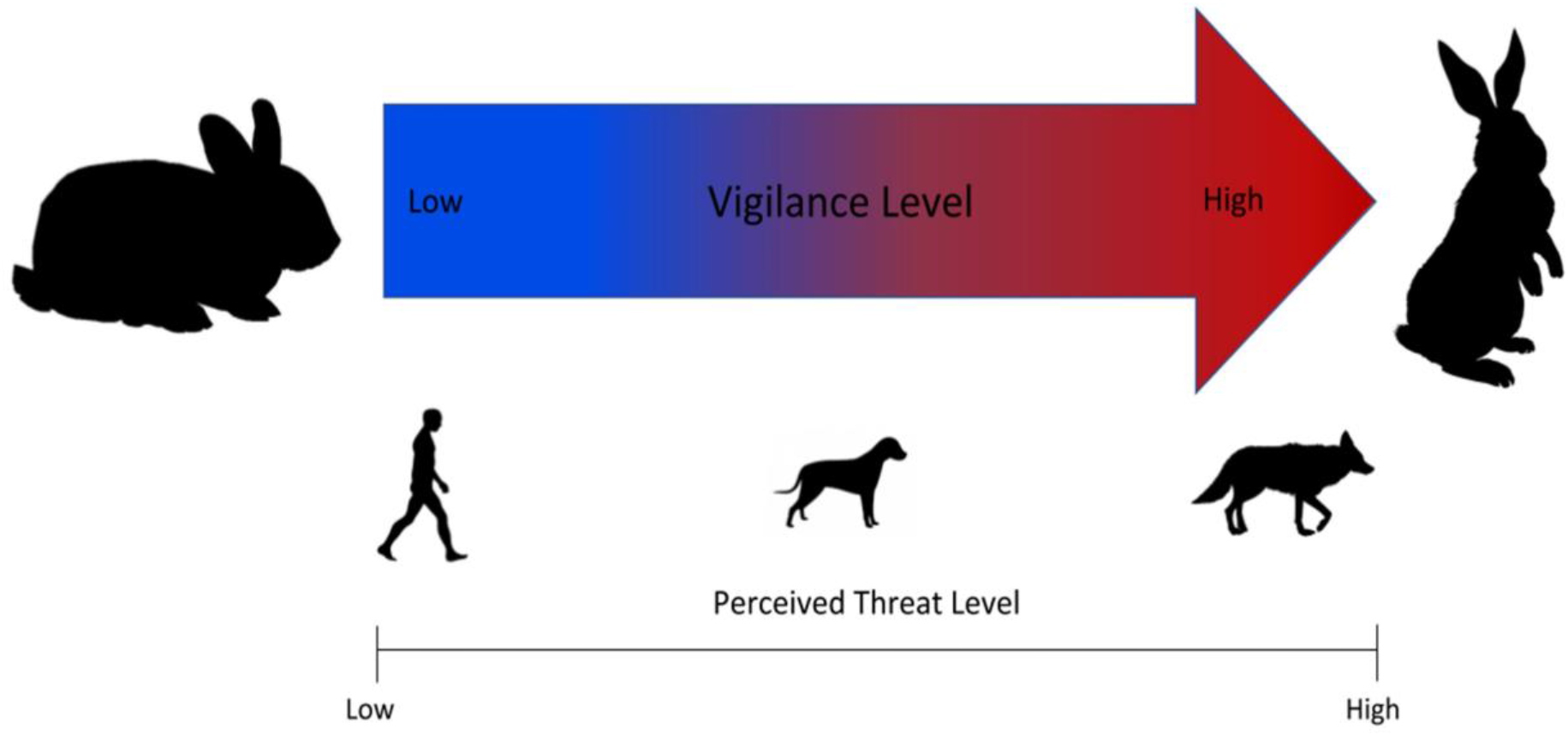
Expected vigilance response of cottontail rabbits with scaled response across natural and anthropogenic threats.

## Materials and Methods

### Study Site

We implemented a systematic camera survey throughout metro parks in Detroit, the largest city in Michigan covering 359.2 km^2^ of land (Figure 2). The declining city holds a human population of 672,000 people with an average density of ~5,144 people per square mile (U.S. Census Bureau, 2016). The Detroit metro park system contributes to the green space and available habitat for wildlife within the city. All 28 total parks sampled within the city are impacted directly or indirectly by humans and are embedded within an urban matrix including roads, neighborhoods, and buildings. The parks range in size from ~0.016 - 4.79 km^2^ with varying levels of vegetation and human influence. In Detroit, the largest native carnivore present is the coyote. However, domestic dogs are also present and may exert pressures on the coyote’s natural prey species such as rabbits.

**Figure 2.**
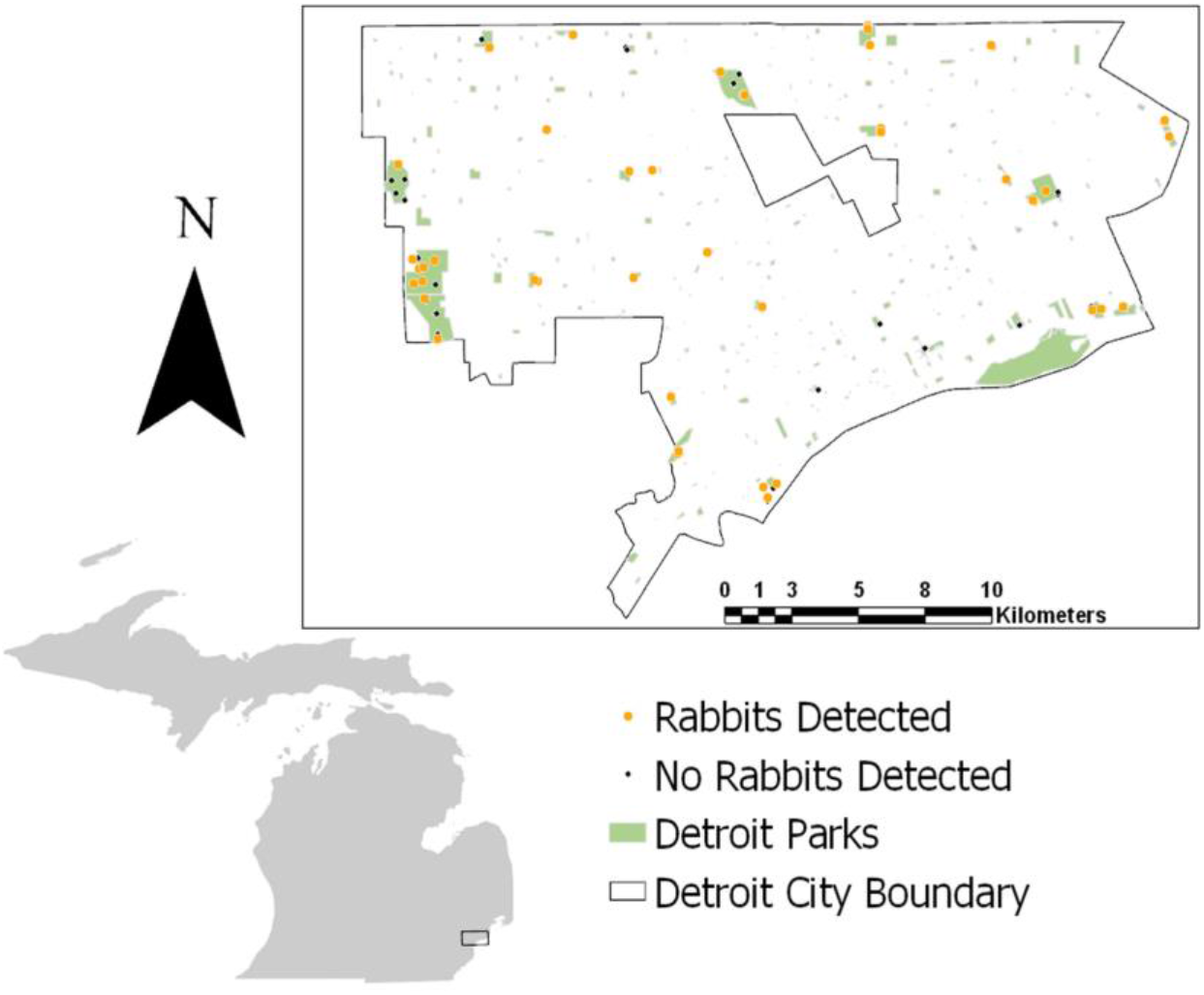
Study site in Detroit with dots indicating camera placement from 2017-2020. Orange dots indicate camera stations where rabbits were detected in at least one of the years of study. Black dots indicate camera stations where no rabbits were detected in any years.

### Data Collection/Camera Survey

We deployed unbaited, remotely triggered cameras (Reconyx© PC 850, 850C, 900, 900C) throughout city parks to monitor the wildlife community from October – March in 2017-2020. Placement within the parks was determined based on evidence of wildlife presence such as scat, and vegetation type. Park size determined the number of cameras deployed, ranging from 1-7 cameras. For parks with multiple cameras, we deployed cameras with a minimum distance of 500m between individual cameras. Cameras were affixed to medium sized trees approximately 0.5-1m off the ground. We programmed cameras to take three images when triggered at high sensitivity with one second between each image and a 15-second quiet period. Every image was independently sorted and confirmed by at least two members of the Applied Wildlife Ecology Lab at the University of Michigan. We only used images confirmed as rabbit as well as their associated threat species of interests: humans, domestic dogs, and coyotes. Both gray (*Urocyon cinereoargenteus*) and red foxes (*Vulpes vulpes*) are also potential predators of cottontails, but sample sizes were too low to include in our analysis. Team members were excluded from human images.

### Hotspot Analysis and Occupancy Modeling

To determine the level of risk from each of our three potential predator focal species, we used two method to capture their spatial variation in parks across Detroit. First, we used kernel density analysis to construct utilization distributions from rabbit, human, coyote, and domestic dog camera triggers in ArcMap (v. 10.6.1). To test for significant spatial clustering (i.e., hotspots), we applied the Getis-Ord-GI* statistic to species triggers, which summarizes spatial autocorrelation with resultant high positive z-scores indicating clustering and low negative z-scores indicating dispersion (Getis and Ord 1992). Specifically, significant trigger hotspots and coldspots are derived from z-scores greater than 1.96 and less than 1.96 (α < 0.05), respectively. Finally, we overlaid significant trigger hotspots for rabbits with associated threats to determine if rabbits avoided hotspots for humans, dogs, or coyotes across the city. In other words, we assessed whether trigger hotspots for rabbits were congruent with any of the threats. Evidence of spatial avoidance may represent a sufficient evasion strategy that necessitates less vigilance behavior.

Second, we constructed single-species, single-season occupancy models for humans, domestic dog, and coyotes, which corrects for imperfect detections from repeated surveys (MacKenzie *et al.* 2003; Mackenzie & Royle 2005). In our case, we used one-week sampling interval to generate detection histories. By holding occupancy constant, we first built detection models with camera model (CAM), understory vegetation at camera (UAC), number of trap nights (TN), and park size (AREA) as covariates. We then used the top detection model to build occupancy models with housing density within 500 meters (HOUSE), prey trap success (PREYTS), UAC, and AREA. PREYTS was calculated at the camera level as the ratio of cottontail rabbit, squirrel, chipmunk (*Tamias striatus*) and small mammal total triggers by number of trap nights. We identified top models using Akaike’s Information Criterion corrected for small sample sizes (AICc) based on the lowest ΔAIC and greatest weight (*w*). We also assessed goodness-of-fit for each model using the chi-squared discrepancy method in the ‘ResourceSelection’ package. We constructed detection histories using ‘camtrapR’, and completed occupancy modeling in ‘unmarked’ packages. All analysis was completed in Program R.

### Vigilance Scoring

We extracted behavioral information from images in order to quantify vigilance response in cottontail rabbits. For each image containing a rabbit, we scored vigilance based upon the position of the body and head (Figure 3). For images with two individuals, each individual was given its own classification and counted as independent from other individuals in the image. Rabbits were considered “vigilant” if their head was in an upright position; while “non-vigilant” was assigned when their head was down in a foraging position. For images where the rabbit did not display an obvious head up or head down stance, we used six other classifications: moving, active, eating, sniffing, out of frame, and unknown. “Moving” included any rabbit in motion, which was often indicated by motion blur in the images. We considered moving to be a potential indicator of vigilance as it could denote rabbits leaving an area due potentially to a detected threat. “Sniffing” included rabbit attention turned to monitoring an aspect of its environment with its head up such as sniffing twigs. Because we are investigating the impact of canid species on rabbit behavior and canids often mark their territory (Bowen and Cowan 1980), we considered sniffing to potentially indicate vigilance as it is a show of risk assessment. Both sniffing and moving were left out of our initial vigilant vs non-vigilant analysis but were included in the vigilant category in our extended analysis. “Active” was used for activity where the animal’s attention was pointed inward at themselves. This included any rabbits scratching, licking, or otherwise attending to their fur, this also included stretching. “Eating” was used in the event that a rabbit had its head up, but clearly had vegetation in its mouth or the image series showed it chewing. Although both active and eating involve attention being pointed inward at the rabbit, we did not include them as non-vigilant in our analysis as we could not confirm non-vigilance. “Out of frame” included any images where the rabbit exited the frame of the picture and nothing was in the image. Images that were sorted as out of frame were removed from the data set and not counted in the final total. Finally, “unknown” was used for rabbits where only parts of the whole body were in the picture, the head was too blurry to determine, or if the body position could not be determined for any other reason. Unknown photos were also removed from the final total. Each individual was only designated one category per each image in which it appeared. All images with rabbits present were used to best estimate the amount of time actually spent in front of the camera at the particular station. We only used photos where rabbits were in the frame, meaning our photos are estimates of time spent in frame. Each image was scored independently for vigilance by at least two members of the Applied Wildlife Ecology Lab at the University of Michigan. Any discrepancies that were not resolved resulted in classifying the image as unknown.

**Figure 3.**
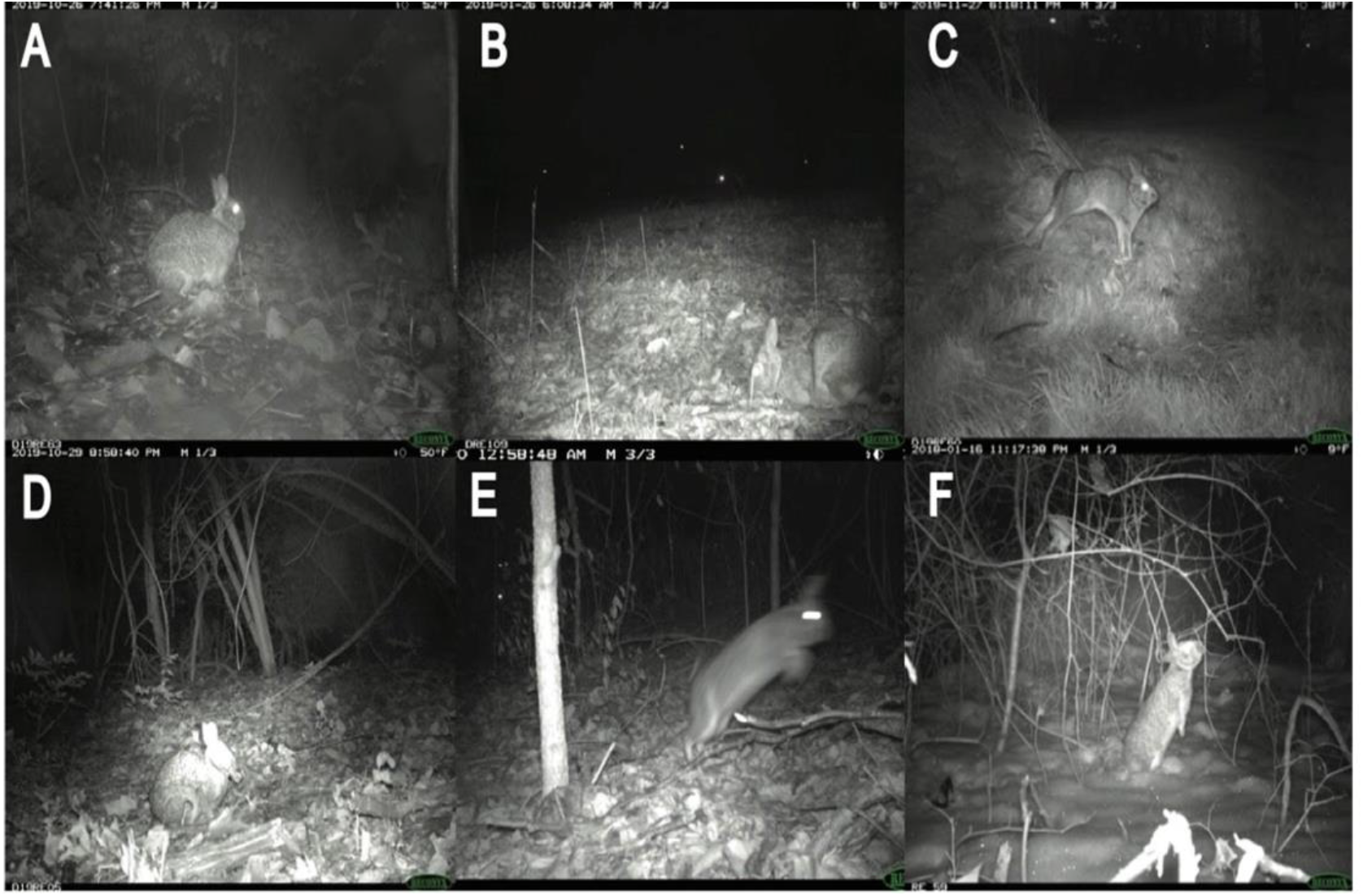
Vigilance classifications based on body position of cottontail rabbits: A) vigilant, head-up; B) non-vigilant, head down; C) active; D) eating; E) moving; and F) sniffing.

We calculated multiple metrics of vigilance as a response variable to each risk factor. Initially, we used the raw number of images classified as vigilant per camera. Our second measure of vigilance was the ratio of vigilant photos to the total number of photos. This was used as a proxy for the relative amount of time spent being vigilant at each camera. For both these metrics, we started with just vigilant and non-vigilant and then expanded the classification of vigilant beyond head up versus head down and included moving and sniffing as vigilant. We used the total raw counts for these combined categories as well as the ratio of those categories out of the total number of detections as our “vigilant” response variable.

### Statistical Analysis

We used negative binomial generalized linear models (GLM.nb) to determine which factors best-explained cottontail rabbit vigilance across cameras. We used results from the hotspot analysis to identify locations of significant high use based on kernel density estimates from detection data to categorize threat levels for humans, domestic dogs, and coyotes. This resulted in a binary explanatory variable indicating whether a hotspot was presence (1) or absence (0) for each threat. We also used occupancy estimates from top models for coyotes (COYO), humans (HUMO), and domestic dogs (DOGO) as threat covariates. We also included environmental and abiotic factors in our analysis. We calculated distance from each camera station to water sources (WATER), to roads (ROADS), and the area of each park (AREA, in acres) using ArcMap. We quantified understory cover (VEG) as a binary variable of whether trees, tall shrubs, bushes, or grasses were present or not in the field of view at the camera level.

Support for models was evaluated using Akaike’s Information Criterion (AICc) to select top-performing model (Δ AIC<2) with highest weight (*w*). We also assessed goodness-of-fit for each model using the chi-squared discrepancy method the ‘ResourceSelection’ package. We completed modeling in the ‘lme4’ package and model selection in the ‘MuMIn’ package. All analysis was completed in Program R.

## Results

We obtained 8,165 cottontail rabbit detections from 58 camera locations in Detroit across 11,616 trap nights from our 2017-2020 surveys (Table 1). The average trap night per camera for the survey period was 99.8 (Range: 18-121) including two cameras which malfunctioned after 18 days, excluding the outliers the average was 101.2 (Range: 74-121). For parks with > 1 camera station, cameras were spaced on average 1.4 km apart within parks spaced an average distance of 3.2 km apart. We recorded 1,345 human detections at 27 camera stations, 484 domestic dog detections at 33 stations, and 271 coyote detections at 29 stations. Three stations (one in 2017 and two in 2019) had no coyote, domestic dog, or human detections. No cameras had significant trigger densities for all three threat species at the same station for the entire duration of study based on Getis-Ord Gi* statistics (Figure 4). Instead, coyotes had significantly high trigger densities to form a hotspot at only one station in 2019. Domestic dogs had hotspots at the same station across two different years. Humans had hotspots at three stations across the three years of study, with two of those stations recurring across years. Rabbits had significant trigger densities at the same two stations across two years. We found spatial aggregation of rabbits with dog at one hotspot location in two years. However, we saw no significant overlap in hotspots between rabbits and humans or coyotes (Figure 4).

**Table 1.** Number of detections for cottontail rabbits and associated threats tested that may influence their vigilance behavior in urban parks, Detroit Michigan 2017-2020.

| **Park** | **Rabbit** | **Human** | **Domestic**<br>**Dog** | **Coyote** |
| --- | --- | --- | --- | --- |
| Balduck | 495 | 22 | 44 | 0 |
| Bishop | 204 | 4 | 9 | 0 |
| Butzel | 2111 | 1 | 5 | 3 |
| Chandler | 18 | 3 | 7 | 7 |
| Comstock | 48 | 144 | 26 | 1 |
| Conner | 288 | 4 | 4 | 21 |
| Eliza Howell | 42 | 0 | 1 | 20 |
| Farwell | 325 | 47 | 31 | 1 |
| Fields | 28 | 25 | 36 | 0 |
| Ford | 259 | 8 | 32 | 6 |
| Fort Wayne | 1552 | 3 | 9 | 16 |
| Hammerberg | 120 | 0 | 2 | 1 |
| Lasky | 12 | 79 | 0 | 0 |
| Maheras | 102 | 7 | 4 | 29 |
| Marruso | 1005 | 21 | 77 | 0 |
| McCabe | 3 | 0 | 0 | 0 |
| O'Hair | 30 | 0 | 8 | 0 |
| Palmer | 42 | 7 | 8 | 8 |
| Patton Memorial | 557 | 28 | 3 | 26 |
| Romanowski | 75 | 7 | 0 | 0 |
| Rouge | 636 | 4 | 22 | 32 |
| Stoepel #2 | 213 | 13 | 29 | 3 |

**Figure 4.**
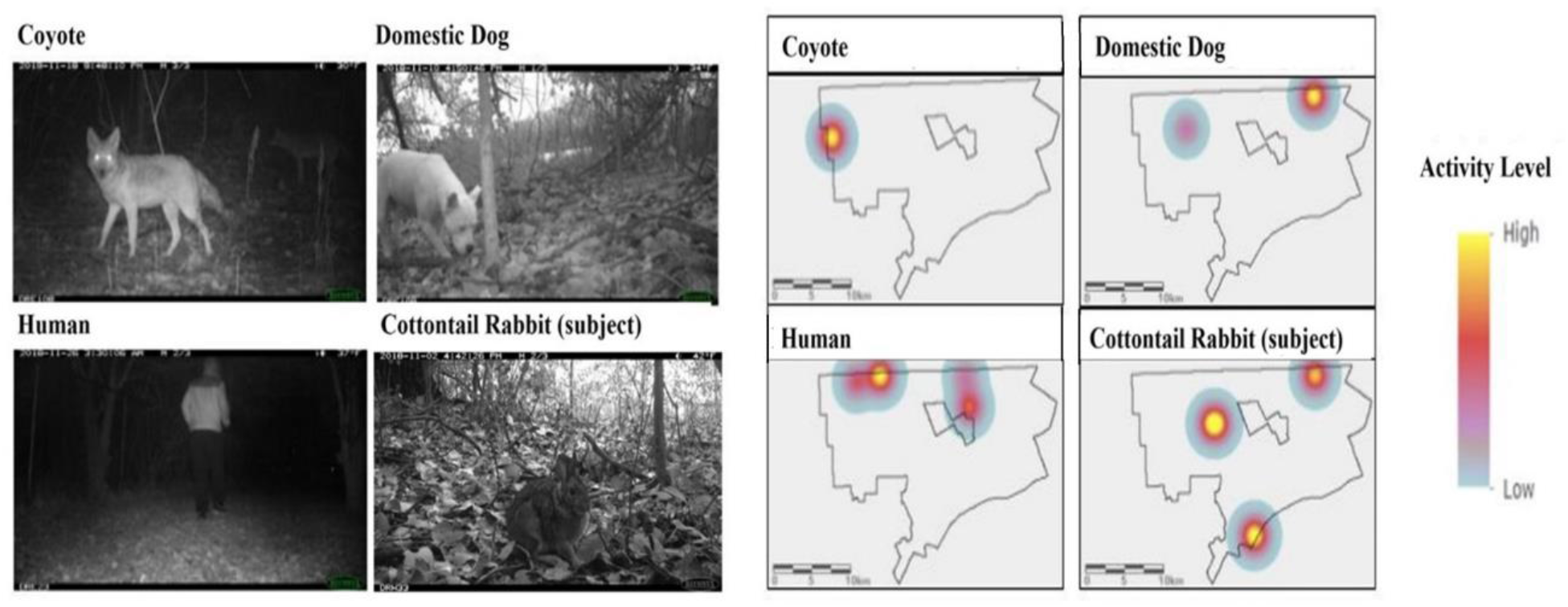
Spatial use within Detroit for rabbits and their three threat species as shown by significant hotspots based on kernel density activity patterns from camera images in the city of Detroit parks from 2017-2020.

Top occupancy models for all threats included HOUSE with PREYTS being important for both canid species (Table S1). Detection models highlighted SIZE for all threats as important as well as TN, UAC, and CAM for humans and domestic dogs. Although comparable, estimates from top models indicated occupancy was highest for humans and lowest for coyotes throughout Detroit city parks (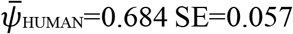; 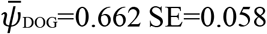; 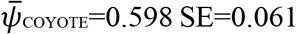).

Of the rabbit detections, with vigilance being determined by head position, we categorized 2,774 images as vigilant (i.e., head-up, 34%) and 1,327 images as non-vigilant (i.e. head down, 16.3%). We classified the remaining 4,064 photos into the following categories: 17.4% moving, 1% active, 1.8% sniffing, and 1% eating. Over a quarter of the total images were either unknown or out of frame, with these categories both being removed from analysis.

Models further support differential effects of threats on rabbit vigilance (Table 2). The top model (highest *w* with Δ AIC<2) indicated that the presence of domestic dog hotspots (β = 2.63,*p=0.002),* coyote occupancy (β = 0.869,*p*=0.013), vegetation cover (β = 0.735,*p* = 0.031) and distance to water (β=0.0001, *p* = 0.078) all positively influenced vigilance, when response represented the number of images with rabbits exhibited vigilance behavior. Though park size, roads, and human occupancy are in other top models, none of these variables had significant beta coefficients in explaining rabbit vigilance. Results of top models were consistent when using the extended categories of vigilance to include counts of moving and sniffing. The intercept-only model was included in top models when using ratio of vigilance photos. Therefore, we did not have sufficient power to investigate whether other variables explained the variation in the proportion of vigilant photos.

**Table 2.**
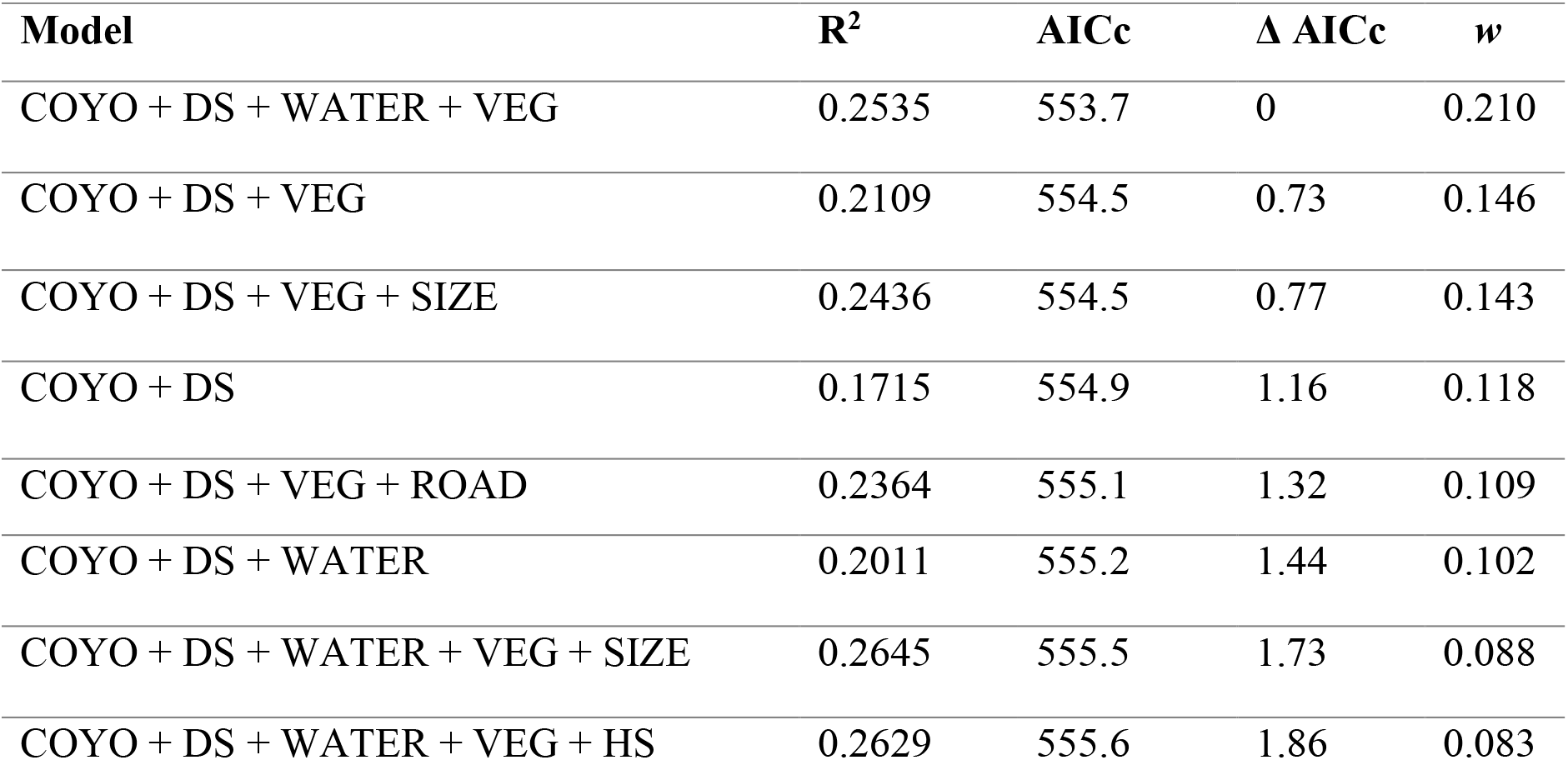
Top models (< 2 Δ AlCc) explained rabbit vigilance behavior using detection data from camera survey in Detroit city parks, 2017-2020. Response variable is number of photos with rabbit head-up. Explanatory variables were: COYO (coyote occupancy), DS (presence of domestic dog hotspot), HS (presence of human hotspot), VEG (vegetation cover), WATER (distance to water), and SIZE (size of the park in acres). Model output for top models includes R^2^, AlCc, Δ AlCc, and model weight (*w*).

## Discussion

Urban wildlife must employ various behavioral strategies to cope with risks in their environment from naturogenic and anthropogenic sources (Blecha et al 2018, Stillfried et al 2017). Like other urban prey species, cottontail rabbits are facing predation threats that are dynamic in an increasingly urbanized world (Santiago-Alarcon 2017, Duarte and Young 2011, Mccleery 2009). We anticipated a scaled response where rabbits showed the lowest vigilance in areas of high human density, then progressively increased with in areas of high domestic dog density and even more in areas of high coyote density. Our analysis showed that occupancy of coyote positively influenced vigilance, consistent with expectations. We did find that rabbit vigilance behavior was heightened in hotspots of domestic dogs across the city. Further, consistent with our expectation, rabbit vigilance was not significantly affected by human occupancy suggesting more acclimation in a human-dominated landscape. Similarly, Gámez and Harris (in press) found no response of human occupancy on carnivore occupancy throughout Detroit in the same parks we surveyed here to access rabbit vigilance behavior. We also found that rabbit vigilance was significantly higher with more vegetation cover, which could be a response to lower visibility to detect predators.

While it is possible rabbits have acclimated to human presence (Dunagun et al. 2019, Samia et al. 2015), their response to domestic dogs indicates that they continue to perceive them as a threat. Domestic dogs are morphologically similar to coyotes, but occupy much higher densities in urban areas and may represent a novel threat similar enough to a natural predator to induce a stronger vigilance response. Coyotes may not occur above the density threshold required to induce behavioral modifications in rabbits in Detroit. Dogs may have functionally replaced coyotes in this capacity posing greater predation risk to cottontail rabbits. Similarly, vigilance behavior increased in association with domestic dogs, but not coyotes in white-tailed deer (*Odocoileus virginianus*) in the mid-Altantic region of the United States (Schuttler et al. 2017). Parsons *et al.* (2016) found that white-tailed deer and gray squirrel avoided humans with and without dogs more strongly than coyotes throughout the southeastern United States. Their findings were notably in contrast with other studies such as Parker and Nilon (2008) that suggested squirrels habituated to human activity in urban areas.

Ziege et al. (2016) found European rabbits (*Oryctolagus cuniculus*) were less vigilant in urban areas as compared to their counterparts in rural areas. This suggests that perhaps the important difference in vigilance lies in the urban-rural gradient, rather than entirely within the urban matrix. Similar to rural areas where there is more vegetation cover than urban areas, we found vigilance increased within areas with more vegetation cover. Rabbits occurring in areas with more vegetative cover increased their vigilance, which could indicate fear that the covered environment may obscure predators. In Missouri, Jones et al. (2014) reported that forest cover did not influence rabbit or squirrel occupancy across an urban-rural gradient study. We also found that as rabbits moved further away from water their vigilance level increased in the urban parks we sampled, which could reflect increased exposure to more disturbed areas in the urban matrix. Urban systems represent a novel landscape for rabbits that requires dynamic changes in vigilance based on the environment and threats of specific locations within the landscape.

Our hotspot analysis indicated very little spatial overlap between species, with domestic dogs and rabbits being the only two species to have significant densities at the same camera location in the same year. As a result, we conclude that generally, rabbits are investing more in spatial avoidance, requiring less effort for vigilance. By mostly avoiding their predators, rabbits may be better able to maintain constant levels of vigilance across the landscape rather than heightening vigilance in areas their predators occupy at significant densities. These hotspots of activity might also be confounded by other factors affecting vigilance that were not incorporated in our models. For example, rabbits might be selecting environments based on proximity to housing, overall vegetation density, or grass cover that might be less desirable for their predators, allowing the rabbits to spend less time being vigilant.

Notably, our analysis was limited in scope by only examining behavior in areas where these species co-occur. It is entirely possible that spatial or temporal partitioning plays a larger role in mediating predator-prey interactions than vigilance solely in prey. We examined interactions within patches in the city, but neglected to examine the amount of interaction occurring between these spaces. Quantifying the level of risks between patches in the city could be the next step in examining threat impacts on prey behavior. Furthermore, seasonality may influence vigilance behavior and interact with food availability (Favreau et al 2018, Périquet et al 2017). Our survey did not sample during warmer months. However, one could argue risk assessment in cottontail rabbits may be more extreme in the winter months when predators are more food-limited.

A growing number of studies on prey behavior have shown increasing evidence for multiple factors affecting predator prey dynamics including human influence and urbanization (Gallo et al 2019, Magle et al 2014). Our work contributes to the growing number of studies on urban wildlife and particularly predator-prey dynamics within urban systems. Further, we underscore that studying behavioral ecology across city topologies including cities where human populations are declining such as Detroit is necessary for understanding how humans, not just their built environment, affect wildlife to better promote coexistence between humans and wildlife (Guerrieri et al. 2012, Herrmann et al. 2016). Understanding the dynamics of predators and their prey in urban systems will be key to the continued coexistence of wildlife and humans in urban spaces. Our results elucidate how a common prey species changes, or fails to change, their vigilance behavior across anthropogenic and naturogenic risk factors in an urban ecosystem. Ultimately, these findings advance our understanding of the adaptability of wildlife in human-dominated environments.

## Author contributions

S.L and N.C.H. conceived the study. S.L. wrote the manuscript and conducted analysis with support from S.G. and N.C.H. S.L. and N.A. designed graphics. All authors contributed to field efforts, data curation, and editing the manuscript. N.C.H. secured funding and supervised work.

## Acknowledgements

First, we recognize implementing our field research with camera traps was conducted on lands originally belonging to the People of the Three Fires indigenous tribal community. We sincerely thank members past and present of the Applied Wildlife Ecology (AWE) Lab at the University of Michigan, specifically R. Malhotra, S. Bower, and G. Gadsden who contributed to the data collection, image sorting, and logistical support of this project. We also thank the University of Michigan Undergraduate Research Opportunities and the honors committee (P. Tucker, A. Ostling) for support of S. Lima during her B.S. degree. Finally, we extend our appreciation to our partners at the Detroit Zoological Society for their financial support and the City of Detroit for collaboration, permits, and access to the parks in our study.

---

**Table S1.**

| Candidate Model | K | AICc | dAICc | w | GOF p-value |
| --- | --- | --- | --- | --- | --- |
| **COYOTE** |  |  |  |  |  |
| $\rho$ (SIZE) $\psi$ (HOUSE + PREYTS) | 5 | 1005.33 | 0 | 0.53 | 0.0799 |
| $\rho$ (.)$\psi$ (HOUSE) | 3 | 1005.82 | 0.48 | 0.41 | 0.0759 |
| **DOG** |  |  |  |  |  |
| $\rho$ (TN +CAM + SIZE +UAC) $\psi$ (HOUSE + UAC + SIZE + PREYTS) | 10 | 1480.97 | 0 | 0.43 | 0.576 |
| $\rho$ (TN) $\psi$ (.) | 3 | 1481.04 | 0.08 | 0.42 | 0.298 |
| $\rho$ (TN) $\psi$ (HOUSE) | 4 | 1483.36 | 2.4 | 0.13 | 0.375 |
| **HUMAN** |  |  |  |  |  |
| $\rho$ (TN +CAM + SIZE +UAC) $\psi$ (HOUSE) | 7 | 1510.98 | 0 | 0.39 | 0.756 |
| $\rho$ (TN+CAM + SIZE +UAC) | 6 | 1512.13 | 1.15 | 0.22 | 0.451 |
| $\rho$ (TN +CAM + SIZE +UAC) $\psi$ (SIZE) | 7 | 1512.22 | 1.24 | 0.21 | 0.215 |
| $\rho$ (TN +CAM + SIZE +UAC) $\psi$ (SIZE + HOUSE) | 8 | 1512.62 | 1.64 | 0.17 | 0.474 |

